# Gel-like inclusions of C-terminal fragments of TDP-43 sequester and inhibit proteasomes in neurons

**DOI:** 10.1101/2021.03.15.435268

**Authors:** Henrick Riemenschneider, Qiang Guo, Jakob Bader, Frédéric Frottin, Daniel Farny, Gernot Kleinberger, Christian Haass, Matthias Mann, F. Ulrich Hartl, Wolfgang Baumeister, Mark S. Hipp, Felix Meissner, Ruben Fernandez-Busnadiego, Dieter Edbauer

## Abstract

TDP-43 inclusions enriched in C-terminal fragments of ~25kDa (“TDP-25”) are associated with neurodegeneration in amyotrophic lateral sclerosis (ALS) and frontotemporal dementia (FTD). Here, we analyzed gain-of-function mechanisms of TDP-25 combining cryo-electron tomography, proteomics and functional assays. TDP-25 inclusions are amorphous with gel-like biophysical properties and sequester proteasomes adopting exclusively substrate-processing conformations. This leads to proteostasis impairment, further enhanced by pathogenic mutations. These findings bolster the importance of proteasome dysfunction in ALS/FTD.

## Introduction

TDP-43 aggregation is the disease-defining pathological hallmark in 90% of patients with amyotrophic lateral sclerosis (ALS) and ~45% of patients with frontotemporal dementia (FTD) (Gao et al, 2017; Prasad et al, 2019). However, we still have a limited understanding of the native structure of TDP-43 inclusions and their role in disease (Gao et al, 2017). Predominantly cytoplasmic neuronal inclusions of TDP-43 correlate strongly with regional neuron loss in spinal cord, motor cortex or frontal-temporal cortical regions (Mackenzie et al, 2013). These inclusions are enriched in 25-35 kDa C-terminal fragments of TDP-43 that contain a glycine-rich low-complexity region (Igaz et al, 2008). Rare autosomal dominant ALS-causing mutations in TDP-43 cluster in this region, although their pathomechanism is still poorly understood, and they have only modest effects *in vitro* and in knockin models (Prasad et al, 2019). However, overexpression of the inclusion-forming 25kDa C-terminal fragment (“TDP-25”) triggers neurodegeneration in mice even without disease-associated mutations (Walker et al, 2015).

Distinct biophysical mechanisms can drive inclusion formation in the context of neurodegenerative diseases: (i) formation of highly insoluble amyloid fibrils with cross β-sheet conformation (Eisenberg & Jucker, 2012), (ii) liquid-liquid phase separation (LLPS) into highly dynamic liquid-like droplets (Gomes & Shorter, 2019), which may solidify and adopt amyloid-like conformations under pathological conditions (Kato et al, 2012; Patel et al, 2015; Qamar et al, 2018). *In vitro,* full length TDP-43 can form liquid droplet through LLPS but short peptide fragments from the C-terminal region form amyloids (Cao et al, 2019; Conicella et al, 2016). By adopting aberrant conformations, aggregated proteins may engage in toxic cellular interactions (Hipp et al, 2019; Olzscha et al, 2011).

Pathogenic mutations in rare familial cases of ALS/FTD are linked to the ubiquitin-proteasome system (UPS) and autophagy, which can clear aggregated and phase-separated proteins, suggesting the proteostasis system is of particular importance at least in genetic ALS/FTD (Deng et al, 2011; Gitcho et al, 2009; Hipp et al, 2019). We have previously shown that poly-GA inclusions in *C9orf72* ALS/FTD disrupt neuronal proteostasis by sequestering proteasomes in a rare transition state, suggesting the cyclic conformational changes required for substrate processing (Collins & Goldberg, 2017) are blocked by the inclusions (Guo et al, 2018).

## TDP-25 forms amorphous inclusions enriched in proteasomes

Here, we aimed to elucidate gain-of-function mechanisms as well as the structure of cytoplasmic TDP-43 aggregates found in sporadic and genetic ALS/FTD focusing on the aggregation-prone TDP-25 fragment (residues 220-414 of full length human TDP-43) (Zhang et al, 2009). Expression of GFP-tagged TDP-25 in rat primary neurons resulted in abundant cytoplasmic inclusions phosphorylated at disease-specific sites (Fig. 1A) (Hasegawa et al, 2008). To determine which pathomechanisms are enhanced by known pathogenic mutations, we additionally used a TDP-25 variant containing eight mutations (G290A, G294V, G298S, A315T, M337V, G348C, N352S, A382T) that individually cause ALS (Prasad et al, 2019). Wild-type and mutant GFP-TDP-25 formed inclusions of similar appearance by light microscopy (Fig. 1A). While GFP-TDP-43 was almost completely soluble in RIPA buffer, a large fraction of wild-type and mutant GFP-TDP-25 was only solubilized upon sequential extraction of the RIPA-insoluble material with 2% SDS (Fig. 1B) indicative of stronger intermolecular interactions in aggregating TDP-25.

**Figure 1:**
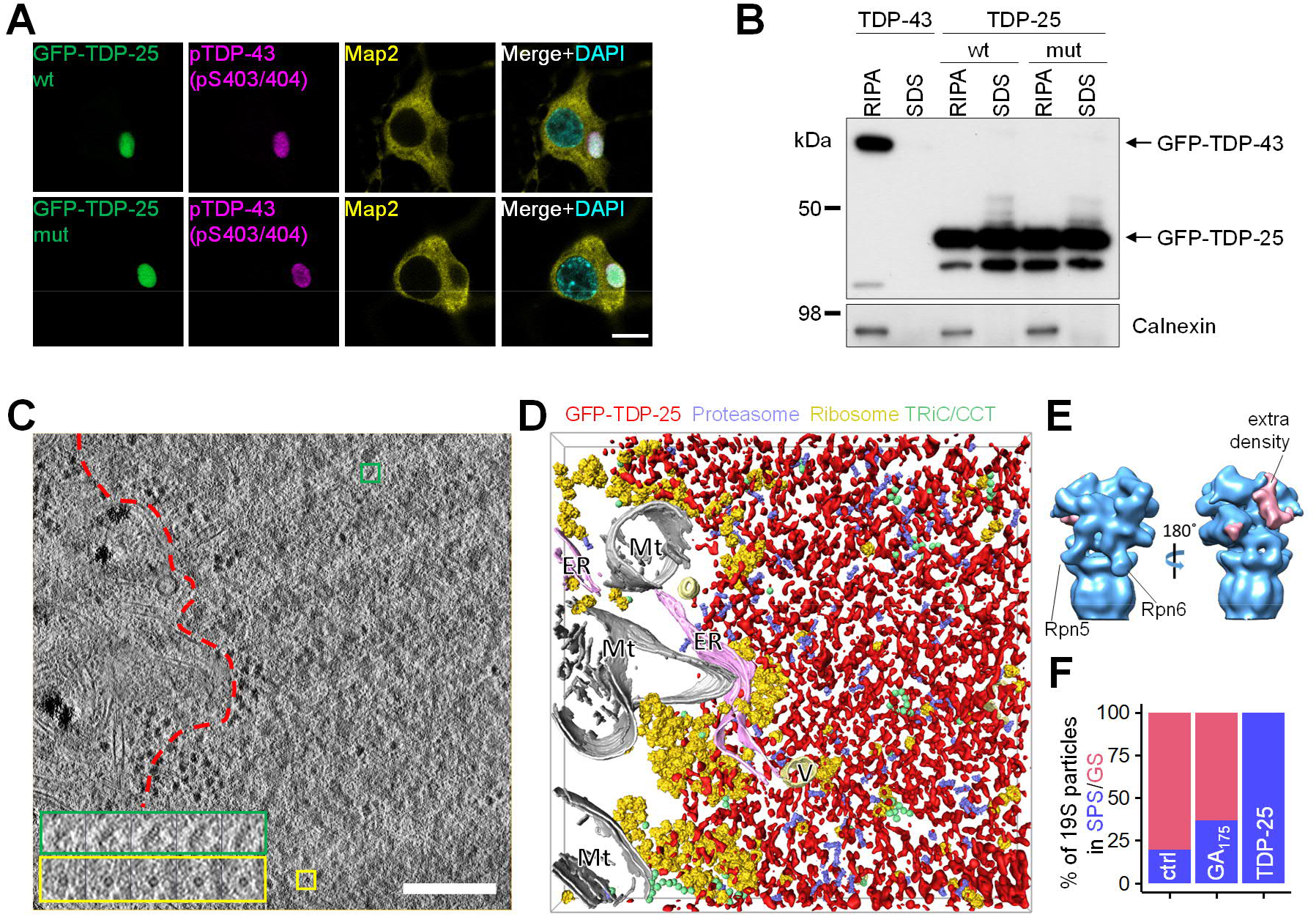
TDP-25 forms amorphous inclusions enriched in proteasomes. A: Primary rat hippocampal neurons were transduced with constructs expressing GFP-tagged TDP-25 (amino acids 220-414 of TDP-43) on day 5 in vitro and cultured for 8 additional days (DIV5+8). Wildtype or variant containing 8 ALS-causing mutations. Immunofluorescence shows disease-related phosphorylation at serine 403/404. Counterstain to label the neuronal cytoskeleton (MAP2) and nuclei (DAPI). Scale bar = 10 μM. B: Hippocampal neurons transduced with GFP-TDP-25 constructs (DIV5+8) or GFP-TDP-43 (DIV5+4 due to higher toxicity) were analyzed by sequential extraction in RIPA buffer followed by 2% SDS. Immunoblots for GFP and loading control Calnexin. C: Tomographic slice of an aggregate within a GFP-TDP-25 wild-type transduced neuron (DIV5+8). Colored boxes show a series of higher magnification tomographic slices of representative protein complexes detected in the tomogram. Green boxes show side views of single-capped, yellow boxes show smaller ring-like structures of 26S proteasomes. Red dotted line segments aggregate area. Scale bar = 200 nm. D: 3D rendering of the aggregate shown in (C). Amorphous aggregate material is labeled in red, proteasomes in violet, ribosomes in yellow, TRiC/CCT chaperones in green, mitochondria (Mt) in white, ER in pink and other vesicles (V) in light yellow. Compare Movie S1. E: Subtomogram averaging of macromolecules in GFP-TDP-25 inclusions reveals the proteasome structure at ~20 Å resolution (see Fig. S1). The positions of Rpn5/PSMD11 and Rpn6/PSDM12 are indicated. Prominent extra densities in the substrate binding region are colored in pink in the 3D rendering. F: Classification based on the conformation of the regulatory particle in TDP-25 inclusions compared to non-transduced control neurons (Asano et al, 2015) and poly-GA inclusions (Guo et al, 2018). GS = Ground State, SPS = Substrate Processing State.

To investigate the downstream consequences of TDP-25 aggregation, we analyzed neuronal GFP-TDP-25 inclusions *in situ* using cryo-electron tomography (cryo-ET), a powerful method to visualize aggregates within their native cellular environment (Bauerlein et al, 2017; Guo et al, 2018). TDP-25 inclusions appeared amorphous and seemingly lacked fibrillar structure, although they were clearly demarcated within the cytoplasm (Fig. 1C/D, Movie S1). These findings are in contrast to poly-Q, poly-GA and α-synuclein neuronal inclusions, which show amyloid-like conformation in neurons in our cryo-ET pipeline (Bauerlein et al, 2017; Guo et al, 2018; Trinkaus et al, 2020). Introducing the eight ALS-causing mutations had no obvious effect on the ultrastructure of TDP-25 inclusion (Fig. S1A).

Similar to our findings in *C9orf72* poly-GA inclusions (Guo et al, 2018), we detected ring-like structures (Fig. 1C) accumulating within TDP-25 inclusions. Sub-tomogram averaging of these rings (Fig. S1B) converged to a proteasome structure at ~20 Å resolution (Fig. S1C). Interestingly, an extra density that was not accounted for by proteasomal subunits was present on the proteasome regulatory particle, possibly reflecting substrates or adaptor proteins (Fig. 1E). Whereas ribosomes were largely excluded from TDP-25 inclusions, we found a ~8-fold enrichment of proteasomes compared to proteasome concentration in control neurons (Asano et al, 2015) (Fig. 1D and S1D). Strikingly, virtually all proteasome particles within the inclusions were in substrate processing states, based on the conformation of 19S regulatory particles (Fig. 1E/F). In comparison, only 20% and 37% of proteasomes were in substrate-processing states in control neurons and poly-GA inclusions, respectively (Asano et al, 2015; Guo et al, 2018) (Fig. 1F). The lack of detectable ground state proteasomes suggests that proteasomes inside TDP-25 inclusions are stalled, as proteasome function requires cyclic transition through activated and ground states (Collins & Goldberg, 2017).

## TDP-25 inclusions are gel-like

Given the amorphous appearance of TDP-25 inclusions, we asked whether they may represent phase-separated liquid droplets. Thus, we analyzed the mobility of GFP-TDP-25 in neuronal inclusions using fluorescence recovery after photobleaching (FRAP) in comparison with known liquid and solid reference proteins, i.e. nucleolar NPM1 and inclusion-forming poly-Q, respectively (Bauerlein et al, 2017; Frottin et al, 2019). In contrast to TDP-25, GFP-NPM1 fluorescence recovered within seconds after bleaching, consistent with high mobility of the protein within the liquid-like nucleolus (Fig. 2A/B). This clearly argues against a liquid droplet character of TDP-25 inclusions. In contrast, nuclear full-length TDP-43 showed a much higher mobile fraction (Fig. S2), in line with its good solubility (Fig. 1B). However, compared to fibrillar poly-Q (huntingtin exon 1 containing 97 glutamines fused to GFP; Htt97Q-GFP), TDP-25 mobility was much higher (~10% vs ~25% recovery at 22 min, Fig. 2C/D). These findings suggest that non-fibrillar TDP-25 adopts a conformation that is best described by a gel-like state. Solidification into hydrogels has been reported for phase separated droplets of FUS, but involves formation of amyloid-like fibrils (Murray et al, 2017; Qamar et al, 2018). A similar process has been postulated for TDP-43, although fibril formation was only detected for much shorter peptides *in vitro* (Cao et al, 2019; Gasset-Rosa et al, 2019; Guenther et al, 2018; Mann et al, 2019). Taken together, GFP-TDP-25 inclusions are clearly less dynamic than classical phase separated compartments, but more dynamic than fibrillar aggregates.

**Figure 2:**
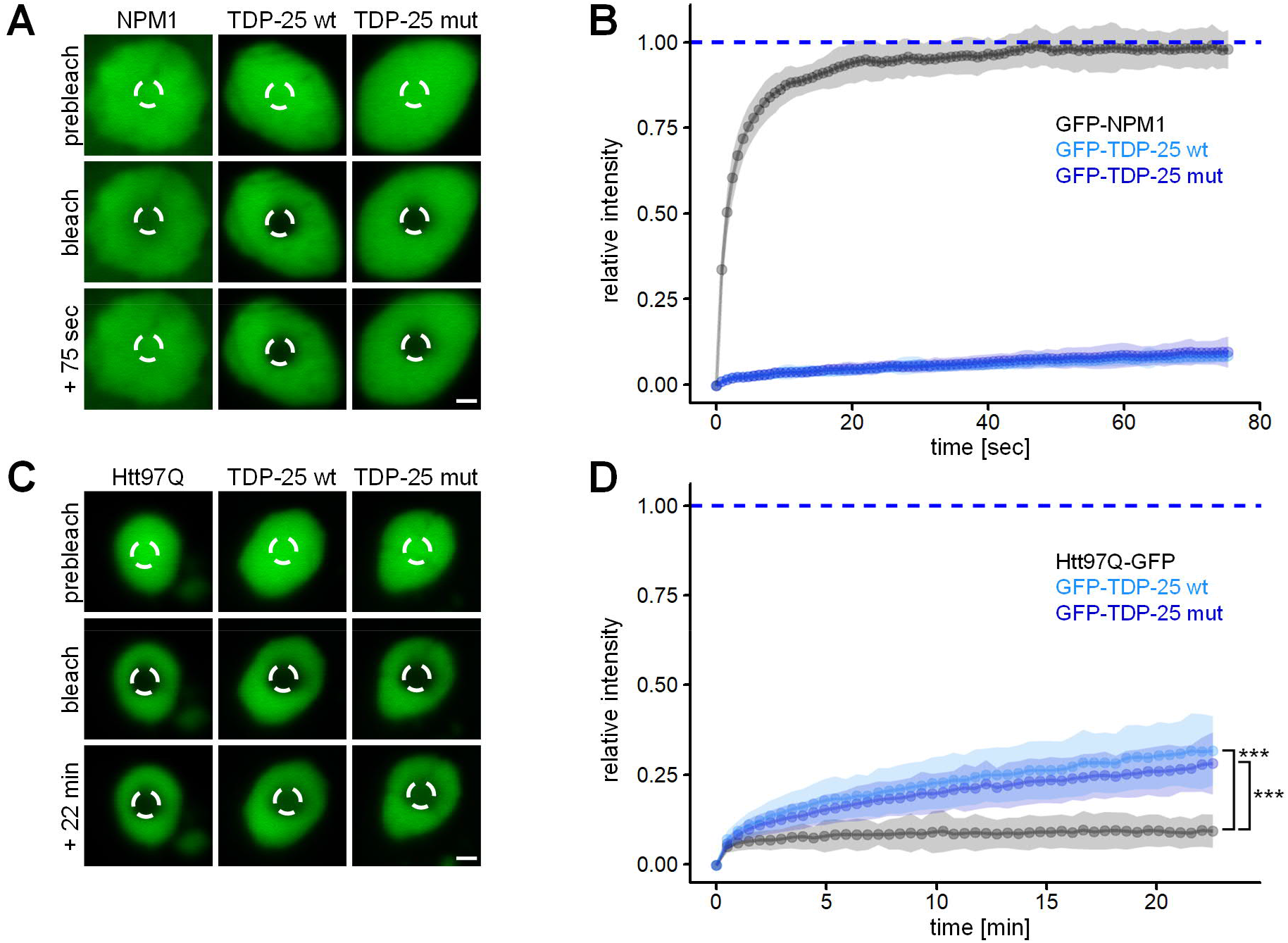
TDP-25 inclusions are neither liquid-like nor solid-like. Primary rat hippocampal neurons were transduced with GFP-TDP-25 variants, GFP-NPM1 or Htt97Q-GFP and analyzed by fluorescence recovery after photobleaching (FRAP) at DIV5+8, except Htt97Q-GFP which was analyzed at DIV5+6 to avoid excessive toxicity. Example images with indicated bleach regions (dashed circles) are shown (A, C). Scale bar = 1.5 μM. Resulting normalized FRAP curves (relative fluorescence intensity over time) with values representing means ± SD are shown in (B, D). (B) GFP-TDP-25 wild-type (n=35 cells), GFP-TDP-25 mutant (n=38 cells), GFP-NPM1 (n=37 cells) out of three independent experiments. In (D), the recovery fraction averaged at >20 min timepoints was compared from four independent experiments. Htt97Q-GFP (n=25 cells) vs GFP-TDP-25 wt (n=23) vs GFP-TDP-25 mut (n=24): H(1)=46.376, df=2, p<0.001, Kruskal-Wallis Test. Htt97Q-GFP vs GFP-TDP-25 wt p=1.9×10^-11^, Htt97Q-GFP vs GFP-TDP-25 mut p=4.3×10^-12^, Pairwise Wilcoxon Rank Sum Tests with Benjamini-Hochberg correction.

## TDP-25 sequesters and inhibits the proteasome

Phase-separated inclusions may form through weak multivalent interactions with other cellular components (Gomes & Shorter, 2019; Martin & Mittag, 2018). To further investigate this gel-like state and the molecular consequences of the TDP-25 inclusions, we analyzed the interactome of TDP-25 and full length TDP-43 using mass-spectrometry-based proteomics. We detected several hundred specific interactors for full length TDP-43 and TDP-25 in primary neurons, in comparison to the GFP-only control (Fig. 3A/B and S3A, Table S1). TDP-43 specific interactors were strongly enriched in splicing-associated proteins consistent with the prominent involvement of TDP-43 in splicing (Fig. 3A/D) (Prasad et al, 2019).

**Figure 3:**
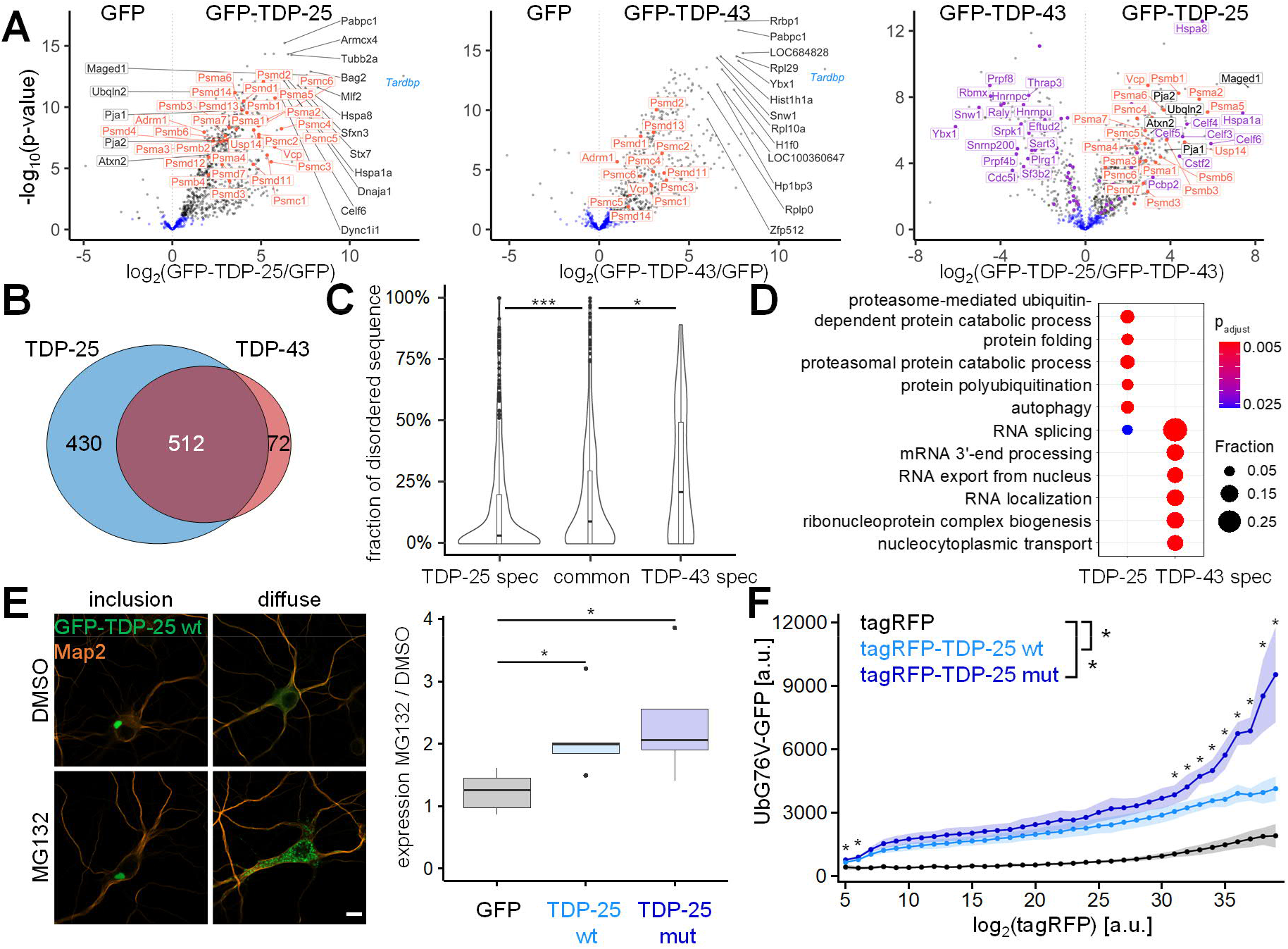
TDP-25 sequesters and inhibits the proteasome. A: Rat primary cortical neurons were transduced with GFP-TDP-25 wild-type, GFP-TDP-43 or GFP at DIV5. Due to high toxicity, GFP-TDP-43 (n=6) and GFP control samples (n=6) were harvested at DIV5+4, while GFP-TDP-25 (n=5) and additional GFP control samples (n=6) were harvested at DIV5+8. Immunoprecipitates were analyzed by LC-MS/MS. Volcano plots indicate enrichment and statistical significance (grey dots: FDR-corrected p<0.05). Proteins associated with GO term “proteasome complex” (GO:0000502) are labeled in orange and proteins associated with GO term “RNA splicing” (GO:0008380) in purple. Full data is list in Table S1. A comparison with previous proteomic studies is shown in Fig. S3A. B: Overlap of GFP-TDP-25 and GFP-TDP-43 interactomes. Significant interactors in FDR-based approach (see methods). C: Combined violin and box plot of the proportion of low-complexity regions (D2P2) of GFP-TDP-25-specific, GFP-TDP-43-specific or shared/common interactors. TDP-25 specific (n=432 Uniprot IDs) vs TDP-43 specific (n=78) vs common interactors (n=481): H(1)=22.276, df=2, p<0.001, Kruskal-Wallis test. TDP-25 specific vs TDP-43 specific: p=0.00054, TDP-25 specific vs common: p=0.00035, TDP-43 specific vs common interactors: p=0.04893, Pairwise Wilcoxon Rank Sum Tests with Benjamini-Hochberg correction. D: Gene ontology (GO) enrichment analysis (biological process) in interactors specific for either GFP-TDP-25 or GFP-TDP-43. Full list of GO terms is found in Table S2. E: Immunofluorescence of hippocampal neurons transduced with GFP, GFP-TDP-25 wild-type or mutant and treated with MG132 (10 μM, 14h, DIV5+8). Representative images of GFP-TDP-25 inclusions and diffuse cytoplasmic staining. Automated quantification of GFP expression normalized to DAPI stained nuclei (not shown) in MG132 treated cells compared to DMSO control. Counterstain to label the neuronal cytoskeleton (MAP2). Scale bar = 10 μM. GFP (n=5 independent experiments) vs GFP-TDP-25 wt (n=5) vs GFP-TDP-25 mut (n=5): H(1)=7.34, df=2, p<0.05, Kruskal-Wallis test. GFP vs GFP-TDP-25 wt: p=0.048, GFP vs GFP-TDP-25 mut: p=0.048, Pairwise Wilcoxon Rank Sum Tests with Benjamini-Hochberg correction. F: A HEK293 cell line expressing the proteostasis reporter UbG76V-GFP was transiently transfected with RFP, RFP-TDP-25 wild-type or mutant. UbG76V-GFP levels were analyzed by flow cytometry 72 h after transfection. The relationship of binned tagRFP fluorescence and UbG76V-GFP fluorescence shows a concentration-dependent accumulation of UbG76V-GFP in cells expressing RFP-TDP-25 wildtype and even more so in the mutant variant. Values shown are means ± SD obtained from n=4 independent experiments. Pairwise comparison using Wilcoxon rank sum test with Benjamini-Hochberg correction was performed for each tagRFP intensity gate respectively. Adjusted p-values for comparison of wild-type and mutant TDP-25 are indicated by asterisks: * denotes p<0.05. tagRFP-TDP-25 wild-type and mutant showed statistically significant differences vs tagRFP control in all gates. Reporter levels were significantly higher for TDP-25 mutant than wild-type in high-expressing cells.

The loss of the N-terminus in TDP-25 led to the loss of 72 interactors compared to full-length TDP-43 but resulted in over 400 new interactions, which may shape biophysical properties and drive gain-of-function toxicity in ALS/FTD (Fig. 3B). To investigate the role of LLPS in inclusion formation in our neuronal model, we analyzed the content of low-complexity domains or intrinsically disordered regions in interacting proteins, as such sequences drive phase separation through multivalent homo-or heterotypic interactions (Martin & Mittag, 2018). Interestingly, the proportion of disordered regions/low-complexity regions was lower in TDP-25 specific interactors than in TDP-43 specific and shared interactors (Fig. 3C), consistent with the higher dynamics of nuclear TDP-43 (Fig. S2). The relative absence of proteins with low-complexity domains within TDP-25 inclusions argues against their formation through LLPS, or may reflect solidification driven by loss of LLPS-promoting low-complexity interactors.

Consistent with our cryo-ET data, proteomics showed that proteasome subunits were highly enriched in the TDP-25 interactome (Fig. 3A/D and Table S2). Furthermore, immunofluorescence confirmed partial sequestration of proteasomes into GFP-TDP-25 inclusions (Fig. S3B), similar to poly-GA inclusions in *C9orf72* ALS/FTD (Guo et al, 2018). Our mass spectrometry data also confirmed the previously reported selective interaction of TDP-25 with other UPS components, such as the ALS-linked Ubiquilin 2 (Cassel & Reitz, 2013; Deng et al, 2011) and the E3 ubiquitin ligases Pja1 (Watabe et al, 2020). These or other UBL domain proteins may contribute to the extra density found on the proteasome regulatory particle (Fig. 1E).

Finally, we investigated the functional consequences of proteasome recruitment to TDP-25 inclusions and asked whether TDP-25 is primarily a substrate or an inhibitor of the proteasome. Treating GFP-TDP-25 expressing neurons with the proteasome inhibitor MG132 increased the level of punctate GFP-TDP-25, without obvious effects on the larger inclusions (Fig. 3E), in line with a previous report (Scotter et al, 2014). Quantification showed a similar 2-fold accumulation of GFP-TDP-25 wild-type and mutant upon proteasome inhibition, suggesting that the proteasome is actively degrading largely soluble GFP-TDP-25 under basal conditions.

Our cryo-ET data suggested alterations in the functional cycle of proteasomes recruited to TDP-25 inclusions. To interrogate whether TDP-25 affects the cellular proteasome capacity, we expressed RFP-TDP-25 in an established UbG76V-GFP reporter HEK293 cell line (Dantuma et al, 2000). We noticed a dose-dependent accumulation of the UPS reporter in cells expressing RFP-TDP-25 wild-type and mutant, indicative of competitive inhibition (Fig. 3F). At high expression levels, mutant TDP-25 impaired overall protein degradation significantly more than wild-type, arguing for a potentially disease-relevant mechanism. These findings corroborate our structural data, indicating that proteasomes enriched in TDP-25 inclusions are functionally impaired.

## Conclusion

Using cryo-ET, proteomics and functional assays in primary neurons, we show that TDP-43 C-terminal fragments adopt a gel-like conformation and disrupt proteostasis by sequestering stalled proteasomes. This gel-like conformation may either reflect a transition state from a liquid compartment to mature aggregates or an alternative pathway to aggregation similar to the physiological cytoplasmic myo-granules of full-length TDP-43 found in regenerating muscle, which are SDS-resistant but lack classical amyloid conformation (Vogler et al, 2018). None of the cytoplasmic inclusions of TDP-25 showed the distinct anisosome-like structure containing a TDP-43 shell and a HSP70 core recently described for RNA-free TDP-43 in the nucleus (Yu et al, 2021), although both TDP-25 and TDP-43 interacted with HSP70 proteins in our model (Hspa1a and Hspa8, see Fig. 3A). Thus, the disease-associated proteolytic fragmentation and cytoplasmic localization of TDP-43 fundamentally change its biophysical properties. Interestingly, so far cryo-ET has only detected proteasome accumulation and impairment within inclusions related to ALS/FTD ((Guo et al, 2018) and this manuscript), but not in other protein aggregates such as poly-Q or α-synuclein (Bauerlein et al, 2017; Schaefer et al, 2019; Trinkaus et al, 2020). Additionally, we show that ALS-causing mutations may further impair proteasome function. These data resonate with previous reports on the crucial role of the UPS for motor neuron function and survival (Bax et al, 2019; Bett et al, 2009; Tashiro et al, 2012), pointing to proteasome dysfunction as a major specific hallmark of ALS/FTD pathogenesis.

## Acknowledgements

We thank Philipp Erdmann, Günter Pfeifer, Jürgen Plitzko and Miroslava Schaffer for electron microscopy support. We thank the Imaging Facility of the Max Planck Institute of Biochemistry. We thank Christian Behrends, Mareike Czuppa, Dorothee Dormann, Bettina Schmid, Ali Rezaei and Qihui Zhou for critical comments to the manuscript. This work was supported by NOMIS foundation (D.E.). M.S.H., F.U.H., Q.G., W.B. and R.F.-B. have received funding from the European Commission (FP7 GA ERC-2012-SyG_318987-ToPAG). D.E., C.H., M.S.H., F.U.H. and R.F.-B. acknowledge funding from the Deutsche Forschungsgemeinschaft (DFG, German Research Foundation) through Germany’s Excellence Strategy - EXC 2067/1-390729940 (R.F.-B.) and EXC 2145 – 390857198 (C.H., F.U.H., M.S.H. and D.E.).

## Methods

### Primary rat neurons

Hippocampi and neocortices from E19 rat embryos were dissected in ice-cold dissection media (HBBS, 1% penicillin/streptomycin, 10 mM HEPES pH 7.3, all from Thermo Fisher Scientific), followed by enzymatic dissociation in dissection media (15 min in 0.15% trypsin for hippocampi, 20 min in 0.25% trypsin, 0.7 mg/ml DNase I for cortices). Hippocampal neurons were plated in Neurobasal media supplemented with 2% B27, 1% penicillin-streptomycin, 0.5 mM L-Glutamine and 12.5 μM L-Glutamate (all Thermo Fisher Scientific) at 85,000 cells/ml on gold grids for cryo-ET experiments, on poly-D-lysine-coated glass coverslips (VWR) in 12-well plates for immunofluorescence, in 35 mm petri dishes with a 14 mm glass bottom inlay (MatTek Corporation) for FRAP analysis and in 6-well plates for biochemical analysis. For mass spectrometry, cortical neurons were plated in 10 cm petri dishes at a density of 400,000 cells/ml in the above media without L-Glutamate supplementation.

### DNA constructs

cDNA encoding for TDP-25 (amino acids 220-414), TDP-43 (amino acids 1-414) and NPM1 were PCR amplified from HEK293FT (Thermo Fisher Scientific) cDNA and subsequently cloned into the FhSynW backbone (human synapsin promoter) with an N-terminal EGFP tag (May et al., 2014). TDP-43 variants with N-terminal tagRFP were cloned in a lentiviral backbone with UbC promoter (FUW3), while human PSMB2 was expressed with a C-terminal tagRFP. TDP-43 point-mutations were introduced using standard PCR methods. The Htt97-EGFP construct was generated by subcloning the open reading frame from Addgene #1186 into FhSynW.

### Lentiviral packaging and transduction

HEK293FT cells of low passage number were seeded into three 10 cm dishes (5×10^6^ cells/dish) using DMEM (Thermo Fisher Scientific) supplemented with 10% fetal bovine serum (Sigma Aldrich), 1% penicillin-streptomycin and 1% Non-Essential Amino Acids (Thermo Fisher Scientific) as described previously (Guo et al., 2018). On the next day, cells were co-transfected with 18.6 μg transfer vector, 11 μg pSPAX2 and 6.4 μg pVSVG using Lipofectamine 2000 (Thermo Fisher Scientific). The transfection media was replaced by plating media supplemented with 13 mg/mL bovine serum albumin (Sigma Aldrich) on the following day. Lentivirus from the cell supernatant was collected 24 h later by ultracentrifugation (87,000 × g 2 h). Finally, lentiviral particles were resuspended in Neurobasal media and stored at −80°C in aliquots.

### Immunofluorescence, Image Acquisition and Quantitative Analysis

Cells grown on coverslips were briefly washed with PBS once before fixing for 10 min at room temperature using 4% paraformaldehyde (Sigma Aldrich) and 4% sucrose (Sigma-Aldrich) in PBS. Primary antibodies (anti-phospho TDP-43 [pS403/404], Cat# TIP-PTD-P05, Cosmo Bio Co., Ltd., 1:1000; anti-Map2, Cat# M1406, clone AP-20, Sigma-Aldrich, 1:250) as well as secondary antibodies (Goat anti-mouse Alexa 555 or Goat anti-mouse Alexa 647, Thermo Fisher Scientific, 1:400) were diluted in GDB buffer containing 0.1% gelatin (Sigma-Aldrich), 0.3% Triton X-100 (Merck), 450 mM NaCl, 16 mM sodium phosphate pH 7.4. Coverslips were incubated with primary antibodies overnight at 4°C and secondary antibodies for 1 h at room temperature, each followed by three washes with PBS. Coverslips were mounted using Vectashield Vibrance with DAPI (Cat# VEC-H-1800, Biozol) to counterstain nuclei.

For colocalization studies, confocal microscopy was performed using an inverted Zeiss LSM800 Axio Observer.Z1 / 7 confocal laser scanning microscope (Carl Zeiss) with a Plan-Apochromat 63×/1.40 Oil DIC M27 objective and equipped with the ZEN 2.5 software package (blue edition, Carl Zeiss). All images were acquired at 2048 × 2048 pixel resolution with 2-times averaging using two GaAsP PMT detectors at 8-bit depth.

To quantify effects of proteasomal inhibition (MG132 vs DMSO), tile scan images (> 100 tiles per condition) were taken on a Leica DMi8 fluorescence microscope using a HC PL APO 40×/0,95 CORR objective, a DFC9000 camera and LAS X Software (Leica Microsystems). Images were quantified via ImageJ/Fiji software (version 1.53c) to determine differences in GFP signal between conditions using a custom script. Integrated Density of GFP-TDP-25 was measured using a fixed threshold and the resulting median was normalized to the number of detected nuclei. DAPI-stained nuclei were counted as particles with circularity factor 0.5-1.0 and size between 60-150 μM^2^ after thresholding and binary water shedding.

### Cellular fractionation and immunoblotting

Transduced neurons were washed with PBS and lysed in RIPA buffer (137 mM NaCl, 20 mM Tris-HCl pH 7.5, 0.1% SDS, 10% glycerol, 1% Triton X-100, 0.5% deoxycholate, 2 mM EDTA) freshly supplemented with 67 U/ml Benzonase Nuclease (Sigma Aldrich), protease inhibitor cocktail (Sigma Aldrich) and phosphatase inhibitor cocktail (Sigma Aldrich) on ice for 30 min. After centrifugation (18,000 × g, 30 min, 4°C), the supernatant was collected as RIPA-soluble fraction. Pellets were resuspended in 2% SDS buffer (2% SDS, 100 mM Tris-HCl pH 7.0) and incubated for 2 h at room temperature. Upon centrifugation (18,000 × g, 30 min, 4°C), the supernatant was collected as SDS-soluble fraction. Samples were denatured for 10 min at 95°C after adding 3× loading buffer (200 mM Tris-HCl pH 6.8, 6% SDS, 20% glycerol, 0.1 g/ml DTT, 0.1 mg Bromophenol Blue). Equivalent fractions of the total cell population were loaded on 10% SDS-PAGE gels, followed by transfer to Immobilon^®^-P PVDF membranes (Merck). Membranes were blocked in 0.2% iBlock (Thermo Fisher Scientific). The following primary antibodies were used: anti-GFP (Cat# 75-131, clone N86/8, UC Davis/NIH Neuromab Facility) and anti-Calnexin (Cat# ADI-SPA-860, Enzo Life Sciences).

### CLEM, cryo-FIB and cryo-ET

CLEM, cryo-FIB/SEM and cryo-TEM tomographic data collection was performed as descripted in detail before (Guo et al, 2018). In brief, EM grids were mounted onto modified Autogrids sample carriers (Rigort et al, 2012) and then transferred into the cryo-stage of an FEI CorrSight microscope. Images of grid and GFP signal were respectively acquired with FEI MAPS software in transmitted light and widefield mode using 5× and 20× lens. The samples were then transferred into a FIB/SEM dualbeam microscope (Quanta 3D FEG, FEI) using a cryo-transfer system (PP3000T, Quorum). Cryo-light microscope and SEM images were correlated in 2D with MAPS 2.1 software (Thermo Fisher; RRID: SCR_018738). Lamellas were prepared using Ga^2+^ ion beam at 30 kV in the regions of GFP signal with final thickness of 100-200 nm.

The grids were then transferred to an FEI Titan Krios transmission electron microscope for tomographic data collection. For the whole procedure, samples were kept at liquid N2 temperature. Tomographic tilt series were recorded with a Gatan K2 Summit direct detector in counting mode. A GIF-quantum energy filter was used with a slit width of 20 eV to remove inelastically scattered electrons. Tilt series were collected dose-symmetrically between −51° to +69° starting from +12° with an increment of 3° and total dose of 110 e^-^/Å^2^ using SerialEM software (Mastronarde, 2005) at a pixel size of 3.52 Å.

### cET Image processing

Image frames were aligned using Motioncor2 (Zheng et al, 2017). IMOD software package (Kremer et al, 1996) was used for tomogram reconstruction: the tilt series were firstly aligned using fiducial-less patch tracking, tomogram were then reconstructed by weighted back projection of the resulting aligned images. Contrast was enhanced by filtering the tomograms using MATLAB script tom_decov (https://github.com/dtegunov/tom_deconv).

MATLAB with TOM toolbox(Nickell et al, 2005) was used as general platform for image processing. For segmentation, tomograms were rescaled with a binning factor of four. The membranes were firstly segmented automatically with TomoSegMemTV (Martinez-Sanchez et al, 2014) using a tensor voting method, and then manually optimized with Amira. Proteasome, ribosome and TRiC were detected using an template matching procedure with PyTOM software (Hrabe et al, 2012), templates were generated by filtering the corresponding structures from previous work (Guo et al, 2018) to 30 Å. The resulting coordinates were used to crop the full size subtomograms from the original tomograms, which were then CTF corrected, classified and refined using RELION (Bharat & Scheres, 2016). In total 1170 proteasome subtomograms were picked from 11 tomograms with dominant TDP-25 aggregates for further analysis.

### FRAP assay

FRAP (fluorescence recovery after photobleaching) experiments were performed on transduced hippocampal neurons on glass bottom dishes at 37°C in HBSS buffer (Thermo Fisher Scientific) supplemented with 20 mM HEPES (Thermo Fisher Scientific) and 4.5 g/L Glucose using an inverted LSM710 Axio Observer.Z1 confocal laser scanning system (Carl Zeiss) equipped with a Plan-Apochromat 63×/1.40 Oil DIC M27 objective, a PMT detector and the ZEN 2011 software (black edition, Carl Zeiss). Images were acquired with two-line averages. Three pre-bleach images were taken. Within the investigated structure (inclusion or nucleolus), a circular region of interest (ROI) of approx. 1.3 μM diameter was photobleached at 100% laser power and fluorescent recovery was monitored over 75 s (image acquisition in 0.8 s intervals), 22 min (30 s intervals) or 450 s (10 s intervals). For analysis, recorded movies were aligned using the *StackReg* plugin of the ImageJ/Fiji software. Afterwards, the mean intensity values of each region of interest (ROI) were monitored over time with ROI1 being the bleached region, ROI2 the whole nucleolus/inclusion and ROI3 the background noise. Values were then analyzed using easyFRAP (version 9.0.1) by applying the “full scale” normalization method (Rapsomaniki et al, 2012).

### Immunoprecipitation

Transduced neurons were lysed in 2% Triton X-100, 750 mM NaCl, 1 mM KH2PO4, freshly supplemented with 67 U/ml Benzonase Nuclease, 1x protease inhibitor cocktail and 1x phosphatase inhibitor cocktail under constant rotation for 45min at 4°C and then centrifuged at low speed (1,000 x g, 5min, 4°C) to remove cellular debris but retain inclusion proteins (Hartmann et al, 2018). Supernatants were incubated with anti-GFP-antibody (Cat# N86/38, clone N86/38, UC Davis/NIH NeuroMab) pre-bound to Protein G Dynabeads (Cat# 10004D, Life Technologies) on a shaking rotator for 3h at 4°C. Beads were subsequently washed three times (150 mM NaCl, 50 mM Tris-HCl, pH 7.5, and 5% Glycerol) and analyzed by mass spectrometry.

### Sample preparation for LC-MS/MS

Beads were resuspended in 155 μl denaturing buffer (8 M urea, 50 mM Tris-HCl pH 7.5, 1 mM dithiothreitol). Proteins were digested off the beads with LysC (220 ng/sample) for 1 h at room temperature while shaking at 1200 rpm. The suspension was diluted 4-fold to lower urea concentration permitting trypsin digestion. Iodoacetamide (final concentration 5 mM) to alkylate cysteines and trypsin (220 ng/sample) were added to digest for another hour before the supernatant was transferred to another tube and digested overnight. The digest was stopped by addition of trifluoroacetic acid to 1% v/v and half of the peptide solution was further processed by desalting chromatography on three disks of C18 material using the STAGE-tip format (Kulak et al, 2014). Briefly, STAGE-tips were washed with 100 μl buffer B (50% v/v acetonitrile, 0.5% v/v acetic acid), conditioned with 100 μl methanol, washed twice with 100 μl buffer A (2% v/v acetonitrile, 0.5% v/v acetic acid), loaded with sample peptides, washed twice with 100 μl buffer A, and subjected to peptide elution by 60 μl of buffer B. The eluate was evaporated to dryness in a vacuum concentrator. Peptides were re-suspended in 10 μl 2% v/v acetonitrile, 0.5% v/v acetic acid, 0.1% v/v trifluoroacetic acid and stored at −20°C. Peptide concentration was measured spectrophotometrically at 280 nm and 500 ng of peptide/sample were subjected to LC-MS/MS analysis.

### MS data acquisition

Peptides were separated on an EASY-nanoLC 1200 HPLC system (Thermo Fisher Scientific) via in-house packed columns (75-μm inner diameter, 50-cm length, and 1.9-μm C18 particles [Dr. Maisch GmbH]) in a gradient of buffer A (0.5% formic acid) to buffer B (80% acetonitrile, 0.5% formic acid). The gradient started at 5% B, increasing to 30% B in 40 minutes, further to 60% B in 4 min, to 95% B in 4 min, staying at 95% B for 4 min, decreasing to 5% B in 4 min and staying at 5% B for 4 min at a flow rate of 300 nl/min and a temperature of 60°C. A Quadrupole Orbitrap mass spectrometer (Q Exactive HF-X; Thermo Fisher Scientific) was directly coupled to the LC via a nano-electrospray source. The Q Exactive HF-x was operated in a data-dependent mode. The survey scan range was set from 300 to 1,650 *m/z,* with a resolution of 60,000 at *m/z* 200. Up to the 12 most abundant isotope patterns with a charge of two to five were isolated and subjected to collision-induced dissociation fragmentation at a normalized collision energy of 27, an isolation window of 1.4 Th, and a MS/MS resolution of 15,000 at *m/z* 200. Dynamic exclusion to minimize re-sequencing was set to 30 s.

### MS raw data processing

To process MS raw files, we employed the MaxQuant software version 1.6.0.15 (Cox & Mann, 2008), searching against the UniProtKB rat FASTA database using canonical and isoform protein sequences downloaded in March 2018. Default search parameters were utilized unless stated differently. In brief, tryptic peptides with a minimum length of 7 amino acids, a maximum mass of 4600 Da, and two miscleavages at maximum were searched. Carbamidomethlyation was set as a fixed modification and methionine oxidation and protein N-terminal acetylation as variable modifications. A maximum of five modifications per peptide was permitted. A false discovery rate (FDR) cutoff of 1% was applied at the peptide and protein level. The search feature “Match between runs,” which allows the transfer of peptide identifications in the absence of MS/MS-based identification after nonlinear retention time alignment was enabled with a maximum retention time window of 0.7 min. Protein abundances were normalized with the MaxLFQ label-free normalization algorithm built into MaxQuant (Cox et al, 2014).

### Bioinformatic data analysis

The dataset was further processed and analyzed in the Perseus environment version 1.6.1.3 (Tyanova et al, 2016). Reverse (decoy) hits, proteins only identified by site, potential contaminants, and proteins not quantified in at least 65% of samples of at least one condition (construct and DIV) were removed. Protein abundances were log2-transformed. Missing values for protein abundances were imputed from a normal distribution around the detection limit with a standard deviation of 0.3 times that of the quantified protein abundance distribution combining all samples and a downshift of 1.8 standard deviations. Volcano plot data were generated with the built-in Perseus tool using a SAM statistic with an s0-paramater of 0.1 to integrate the effect size and a permutation-based FDR control set to 5% (Tusher et al, 2001). Proteins significantly (q-value < 5%) associated with TDP-25 or TDP-43 were termed interactors and included in the further analysis.

To analyze the content of low-complexity regions of neuronal TDP-25/TDP-43 interactors, we automatically queried the D2P2 database based on MaxQuant reported UniProt identifier, using the first entry if multiple identifiers were reported in protein groups. Low complexity regions were defined as regions with a consensus of at least 6 of 9 analysis pipelines within D2P2 (Oates et al, 2013). Gene ontology (GO) analysis was performed focusing on the “biological process” categories of differentially immunoprecipitated proteins using clusterProfiler version 3.12 (Yu et al, 2012). A manual selection is shown to best represent the key pathways. The full list of significant GO terms is shown in Tables S2. The MS proteomics data have been deposited to the ProteomeXchange Consortium via the PRIDE partner repository with the dataset identifier PXD024358.

### Flow Cytometry

Flow cytometry experiments were performed as described before (Guo et al, 2018). UbG76V-GFP proteasome reporter cells were dissociated 72h after transfection and analyzed with a Thermo Fisher Attune NxT Flow Cytometer (Thermo Fisher Scientific). To compensate for crosstalk between GFP and tagRFP, HEK293 were transfected with tagRFP constructs only. Raw flow cytometry data were analyzed using FlowJo software (version 9.9, Treestar). The tagRFP channel was subdivided into different gates, corresponding to the log of fluorescence intensity of the transfected protein.

### Data Visualization and Statistical Analysis

Data were visualized and statistically analyzed using R (version 3.6.1). Assumptions for parametric tests were analyzed with Shapiro-Wilk test and Levene’s test. Kruskal-Wallis test and subsequent Pairwise Wilcoxon Rank Sum Tests with Benjamini-Hochberg correction were used as nonparametric tests. Data are plotted as mean ± standard deviation (SD) or box plots/violin plots. In the legends mean ± 95% CI is noted.

---

**Figure S1:**
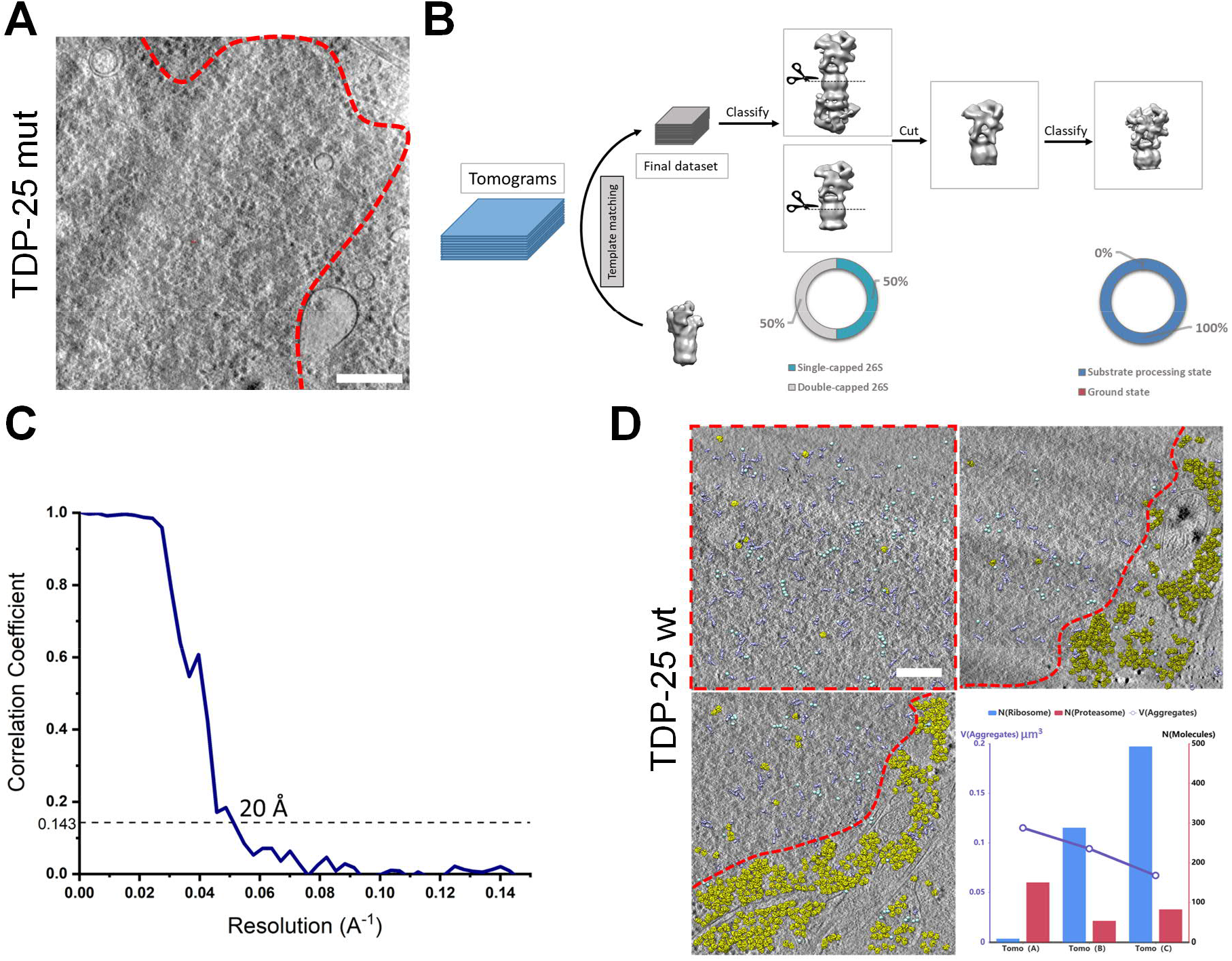
Proteasomes are enriched in TDP-25 inclusions. A: Tomographic slice of an aggregate within a GFP-TDP-25 mutant transduced neuron (DIV5+8). Red dotted line segments aggregate area. Scale bar = 200 nm. B: Workflow of subtomogram averaging and classification. Subtomograms were identified using a low-resolution single-capped proteasome as template. All the proteasomes were firstly classified into single-capped or double-capped. To further analyze the conformation of the regulatory particles, all the proteasomes were cut *in silico* between the β-rings of the core particle, resulting in two independent particles for double-capped ones. Cut out regulatory particles were merged and subjected to a further round of classification. C: Gold-standard Fourier shell correlation curve of the proteasome structure show a resolution of 20Å. D: Molecular mapping in three samples of GFP-TDP-25 wild-type inclusions in transduced neurons (DIV5+8). Regions containing GFP-TDP-25 are outlined in red. For the whole tomogram, proteasomes (purple), TRiC (cyan) and ribosomes (yellow) are mapped to their original positions and orientations using the information from template matching and subtomogram averaging. The numbers of proteasomes and ribosomes detected in the tomograms plotted versus the volume of the region containing GFP-TDP-25 inclusions. Scale bar = 200 nm.

**Figure S2:**
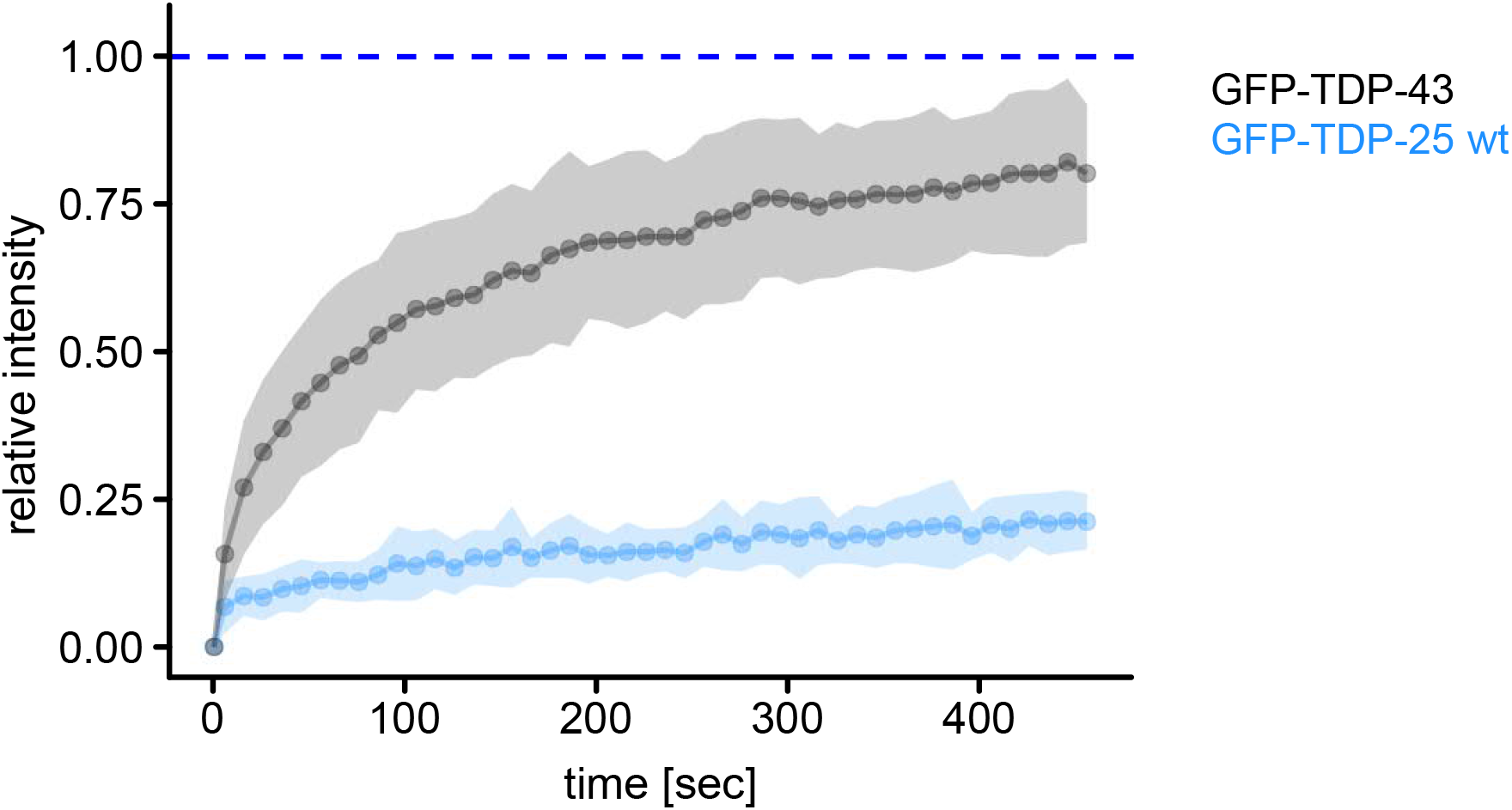
TDP-25 is less mobile than TDP-43. Rat primary hippocampal neurons were transduced with GFP-TDP-25 (DIV5+8) or GFP-TDP-43 (DIV9+4) and analyzed by FRAP. Resulting normalized FRAP curves (relative fluorescence intensity over time) with values representing means ± SD are shown. GFP-TDP-25 wild-type (n=18 cells), GFP-TDP-43 (n=20 cells) from three independent experiments.

**Figure S3:**
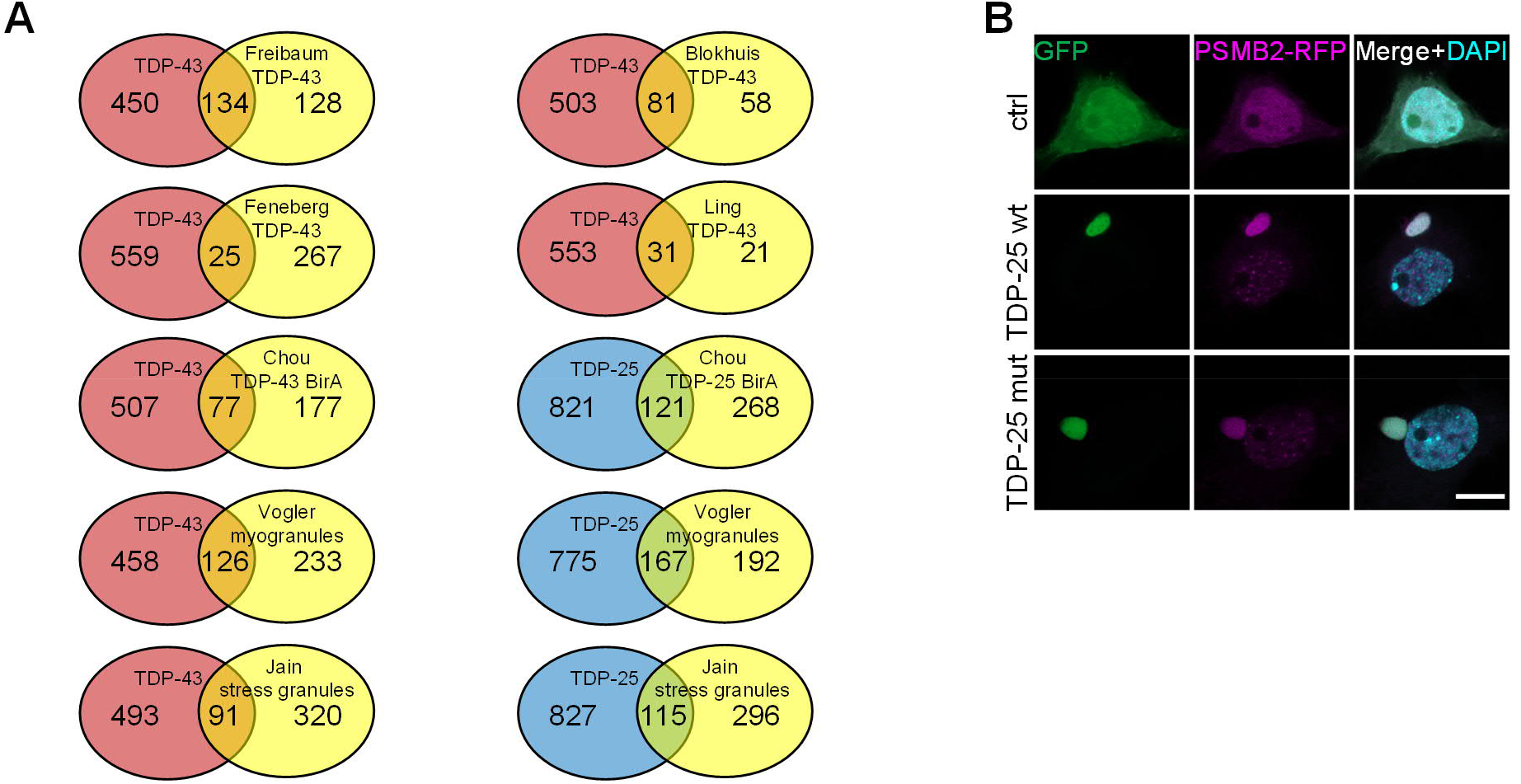
Overlap of TDP-43 and TDP-25 interactome with published datasets. A: Proteins identified in the GFP-TDP-43 interactome (uniquely or in a cluster) were compared to published datasets (Blokhuis et al, 2016; Chou et al, 2018; Feneberg et al, 2020; Freibaum et al, 2010; Ling et al, 2010). LOC and RGD gene names that usually represent minor components of the identified clusters were removed, because such entries could not be mapped based on gene names between species. A core set of 32 proteins from our TDP-43 interactome were found in four (DDX3X, DHX9, ELAVL1, HNRNPA0, HNRNPD, HNRNPH1, HNRNPL, HNRNPM, HNRNPR, ILF2, MATR3, PABPC4, RALY, YBX1) or three (DDX5, DDX6, EFTUD2, EIF3A, HNRNPA3, HNRNPAB, HNRNPC, HNRNPK, HNRNPU, HNRNPUL2, ILF3, NONO, NOP56, NOP58, PABPC1, SNRNP200, SNRPA1, SSB) interactome datasets (Blokhuis et al, 2016; Feneberg et al, 2020; Freibaum et al, 2010; Ling et al, 2010). There was also substantial overlap with a BirA proximity labeling dataset for TDP-43 and TDP-25 (Chou et al, 2018), the stress granule proteome (Jain et al, 2016) and TDP-43 myogranules in regenerating muscle (Vogler et al, 2018). B: Primary rat hippocampal neurons were co-transduced with lentivirus encoding for GFP-tagged TDP-25 variants or GFP and PSMB2-tagRFP lentivirus on day 5. Immunofluorescence images were taken 8 days after transduction (DIV5+8). Counterstain to label nuclei (DAPI). Scale bar = 10 μM.

**Movie S1. Mapping Macromolecules to Poly-GA Inclusions, Related to Figure 2**

Tomographic volume and 3D rendering of the neuronal GFP-TDP-25 inclusion depicted in Fig 1C/D. GFP-TDP-25 (red), 26S proteasomes (violet), ribosomes (yellow), TRiC/CCT chaperonins (green), mitochondria in white, ER in pink and other vesicles in light yellow. The macromolecules are mapped to their original positions and orientations, computationally determined by template matching and subtomogram averaging. Proteasomes are enriched within the GFP-TDP-25 inclusions, while ribosomes are mostly found in its periphery.

## Notes

### Competing Interest Statement

The authors have declared no competing interest.

